# LOXL2 alleviates post-traumatic knee osteoarthritis and pain

**DOI:** 10.1101/2025.05.12.653521

**Authors:** Faiza Ali, Rajnikant Dilip Raut, Chumki Choudhury, Amit Kumar Chakraborty, Cheyleann Del Valle-Ponce De Leon, Pushkar Mehra, Manish V. Bais

## Abstract

Cartilage has limited potential for self-regeneration, and damage can lead to structural, molecular, and functional aberrations, leading to osteoarthritis (OA). Traumatic knee injuries can also lead to cartilage degeneration and post-traumatic OA (PTOA). This study aimed to explore whether lysyl oxidase-like 2 (LOXL2) deletion promotes PTOA-induced transcriptional changes similar to those in human OA, as well as the upregulation of collagen degradation, inflammation, and pain-related gene networks. LOXL2 was found to be downregulated in mouse knee PTOA. Aggrecan promotes specific deletion of *Loxl2* in knee cartilage, shows OA-like molecular changes, and aggravates mouse PTOA. Furthermore, transcriptional analysis revealed the upregulation of cartilage degeneration factors, signatures of inflammatory M1 macrophages, and pain. These *Loxl2* deleted PTOA mice have a molecular resemblance to the human knee OA pathogenic gene signature, which could lead to OA and pain. Interestingly, intra-articular injection of adenovirus-delivered LOXL2 rescued knee joint function, alleviated cartilage degeneration, restored treadmill-running capabilities, and reduced mechanical allodynia by relieving knee joint disability and pain. Taken together, LOXL2 deletion in mice knee promotes the severity PTOA, similar to human OA, implying its potential as a therapeutic candidate for human PTOA.

## Introduction

Osteoarthritis (OA) can develop gradually over a period of years. Trauma, inflammation, and other risk factors degrade the cartilage structure and function in the knee joint. More than six million people visit hospitals because of cartilage loss or abnormalities. The projected lifetime risk of knee OA is 14% ^1^, with annual healthcare costs in the United States reaching $185 billion ^2^. The prevalence is rising with no Food and Drug Administration-approved treatment^3^.

Our study is the first to reveal that lysyl oxidase-like-2 (LOXL2) promotes cartilage regeneration and protects against OA ^4–6^. LOXL2-induced collagen crosslinking enhances the tensile strength of articular cartilage and resistance to collagen proteolysis ^7^. LOXL2 overexpression in human OA articular chondrocytes induces aggrecan (*ACAN*) expression while attenuating the expression of *MMP13* and *NF-kB* signaling ^5^. LOXL2-overexpressing transgenic mice are protected against chemically induced degenerative cartilage changes in the knee joint, restored treadmill running capability, and reduced allodynia compared to their wild-type littermates ^6^. Adenoviral LOXL2 injection protects against progressive cartilage damage in chondrodysplasia (Cho+/™) mice ^6^. However, the role of LOXL2 in traumatic knee injury-induced OA remains unclear and may be crucial for the treatment of human post-traumatic osteoarthritis (PTOA) treatment.

Destabilization of the medial meniscus (DMM) is a common small-animal traumatic knee injury model used to study knee osteoarthritis (OA). It can replicate the progressive cartilage deterioration and joint diseases observed in human OA and PTOA. The DMM model, induced by the surgical transection of the medial meniscotibial ligament, allows researchers to investigate OA’s mechanical and genetic drivers of OA. This model is particularly advantageous for studying pain mechanisms and assessing treatment approaches for OA ^8,9^.

During OA development, M1 macrophages increase in the synovium and circulation ^10^. OA development is strongly linked to synovitis, and synovial macrophages play a key role in synovitis ^11^. M1 macrophages are an activated macrophage subtype that contributes significantly to synovitis, a disorder marked by inflammation in the synovial lining of joints, mainly of proinflammatory cytokines that lead to joint degradation ^11,12^. CD86, TLR2, TLR4, and TNF are key phenotypic markers of proinflammatory M1 macrophages ^12^. Hence, it is necessary to control the levels of proinflammatory macrophages during traumatic injury and knee OA.

Here, we performed DMM or sham surgeries on cartilage-specific *Loxl2* knockout mice to evaluate the efficiency of LOXL2 in controlling the progressive severity of the inflammatory immune response and PTOA development based on the OARSI score. Initially, we assessed the levels of LOXL2, ACAN, and MMP13 in *Loxl2*-deleted mouse knees before DMM and sham surgery using immunohistochemistry (IHC). Both surgeries were performed and the molecular signatures of cartilage degeneration, inflammation, chondrocyte dedifferentiation, and pain sensation were determined using IHC, RNA-seq, and RT-qPCR. Finally, adenoviral LOXL2 was administered intra-articularly in both surgery groups, and recovery from OA and pain was evaluated using the mechanical allodynia test (von Frey filaments), treadmill exhaustion test, and gait analysis to assess stride length.

## Results

### LOXL2 is downregulated in PTOA and *Loxl2 knocoout mice* knee cartilage inhibits ACAN and increase MMP13

To evaluate LOXL2 expression in PTOA, DMM mouse cartilage was compared to that in sham surgery, which showed that LOXL2 expression was downregulated in the knee joint cartilage (Fig. 1a). To evaluate LOXL2 function, we generated tamoxifen (Tm)-inducible, cartilage-specific *Loxl2* knockout mice as illustrated (Fig. 1b) and described in the Methods section. Briefly, *Loxl2* floxed (fl) mice, in which exon 2 was flanked by *loxP* sites, were obtained from Dr. Cano’s laboratory in Madrid, Spain ^13^. These mice were crossed with *Agc1^tm(IRES-CreERT2)^* murine line (Jackson Laboratories, #019148) to generate Tm-inducible Acan-Cre^ERT2^;*Loxl2*^fl/fl^ mice^14–16^. Four-month-old Acan-Cre; *Loxl2*^fl/fl^ homozygous mice (n=8/group, m/f) were injected with either Tm to generate *Loxl2* knockout mice (*Loxl2*^-/-^) or vehicle to generate control mice (*Loxl2*^WT/WT^). IHC confirmed the successful knockout of *Loxl2* in mouse knee cartilage four months post-injection of Tm (Fig. 1c). *Loxl2* deletion decreased ACAN levels (Fig. 1d). Interestingly, the major collagen degradation factor MMP13 was significantly elevated (Fig. 1e). This indicates that LOXL2 is not only a collagen crosslinking factor but also regulates extracelular matrix (ECM) and proteins for normal molecular and cellular homeostasis of cartilage.

**Fig. 1:**
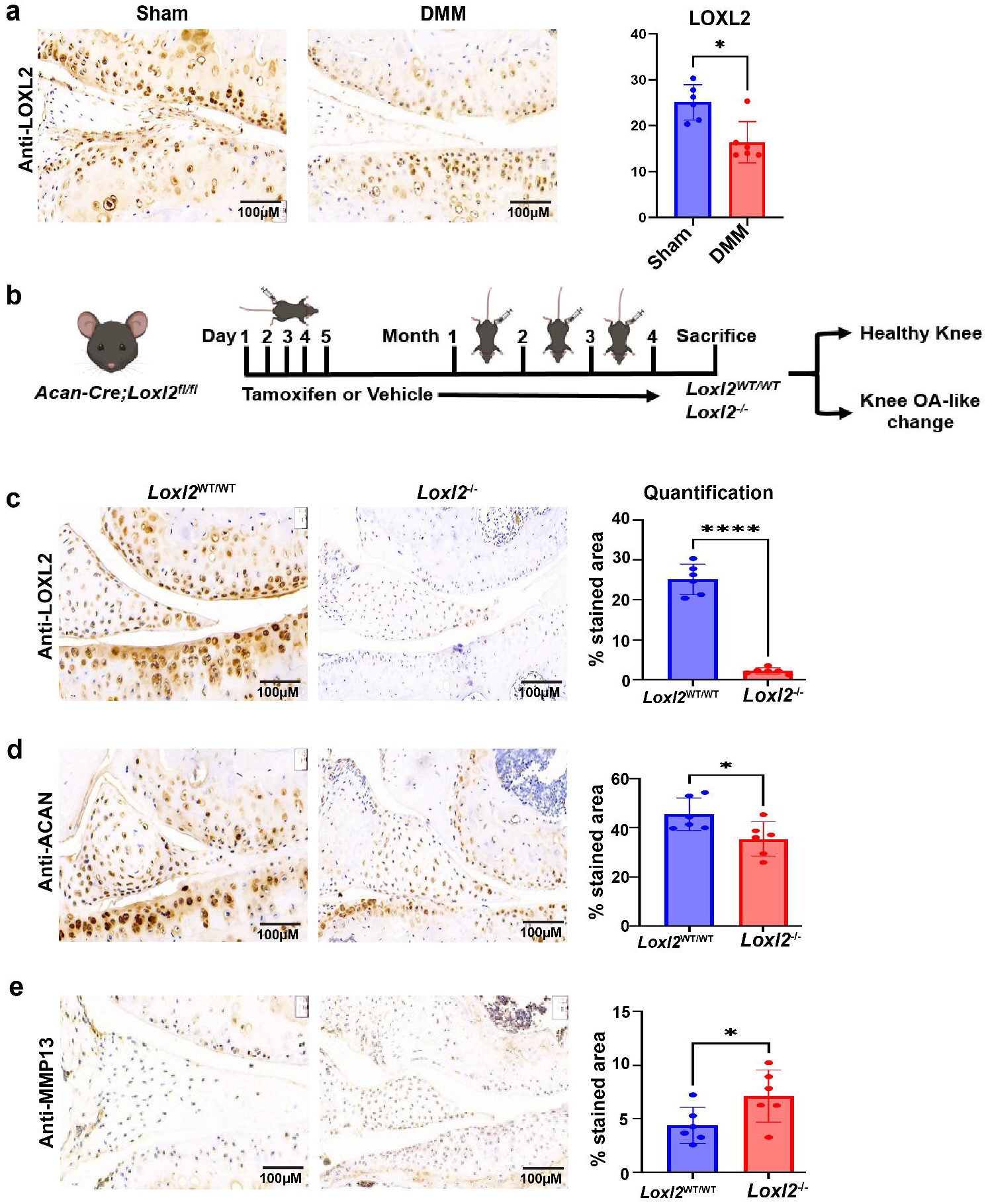
LOXL2 downregulated in DMM-induced mice knee-OA model and *Loxl2* knockout affects ACAN and MMP13. **a)** Expression of LOXL2 after 4 months in mice knee subjected to sham and DMM surgery. **b)** Illustration of cartilage-specific *Loxl2* knockout mice generation. Tamoxifen or vehicle injections were given for five consecutive days to the Acan-Cre;*Loxl2*^fl/fl^ mice followed by single dose every month for four months. c) IHC and bar plot representation of *Loxl2* deletion from mice knee joint. d) Reduced levels of ACAN in mice knee joint with *Loxl2* deletion. e) Increased levels of MMP13 with *Loxl2* deletion. *p<0.05, **p<0.01, ***p<0.001, ****p<0.0001. P values were calculated using *the t test*.

### DMM surgery in *Loxl2* deleted mice increases the severity of knee cartilage damage

To evaluate whether the severity of cartilage damage is increased by injury, we performed Destabilization of Medial Meniscus (DMM) surgeries on the knee of mice to develop a post-traumatic knee-OA (PTOA) model, recapitulating the pathological changes in human PTOA. Specifically, six-month-old *Loxl2*^WT/WT^ (vehicle-injected) and *Loxl2*^-/-^ (Tm-injected) mice were subjected to DMM surgery (n=8/group, m/f). After four months, sectioning, IHC, and high-throughput sequencing analyses were performed. The results showed that DMM-induced changes were more severe in *Loxl2*^-/-^ mice with loss of articular cartilage (Fig. 2a) and higher OARSI scores than those in *Loxl2*^WT/WT^ mice (Fig. 2b). *Loxl2* deletion reduced overall proteoglycan levels according to safranin-O staining; however, the effect was pronounced in the DMM surgery group (Fig. 2a, c). As expected, the *Loxl2* deleted DMM group showed a dramatic reduction in ACAN levels compared with the vehicle-treated group. The Loxl2 deleted DMM group showed a dramatic decrease in ACAN levels compared with the vehicle-treated group (Fig. 2D). Notably, *Loxl2* deletion followed by DMM surgery resulted in increased MMP13 levels compared with the vehicle control (Fig. 2e). These data suggests that *Loxl2* deletion increases the severity of cartilage damage, leading to OA development.

**Fig. 2:**
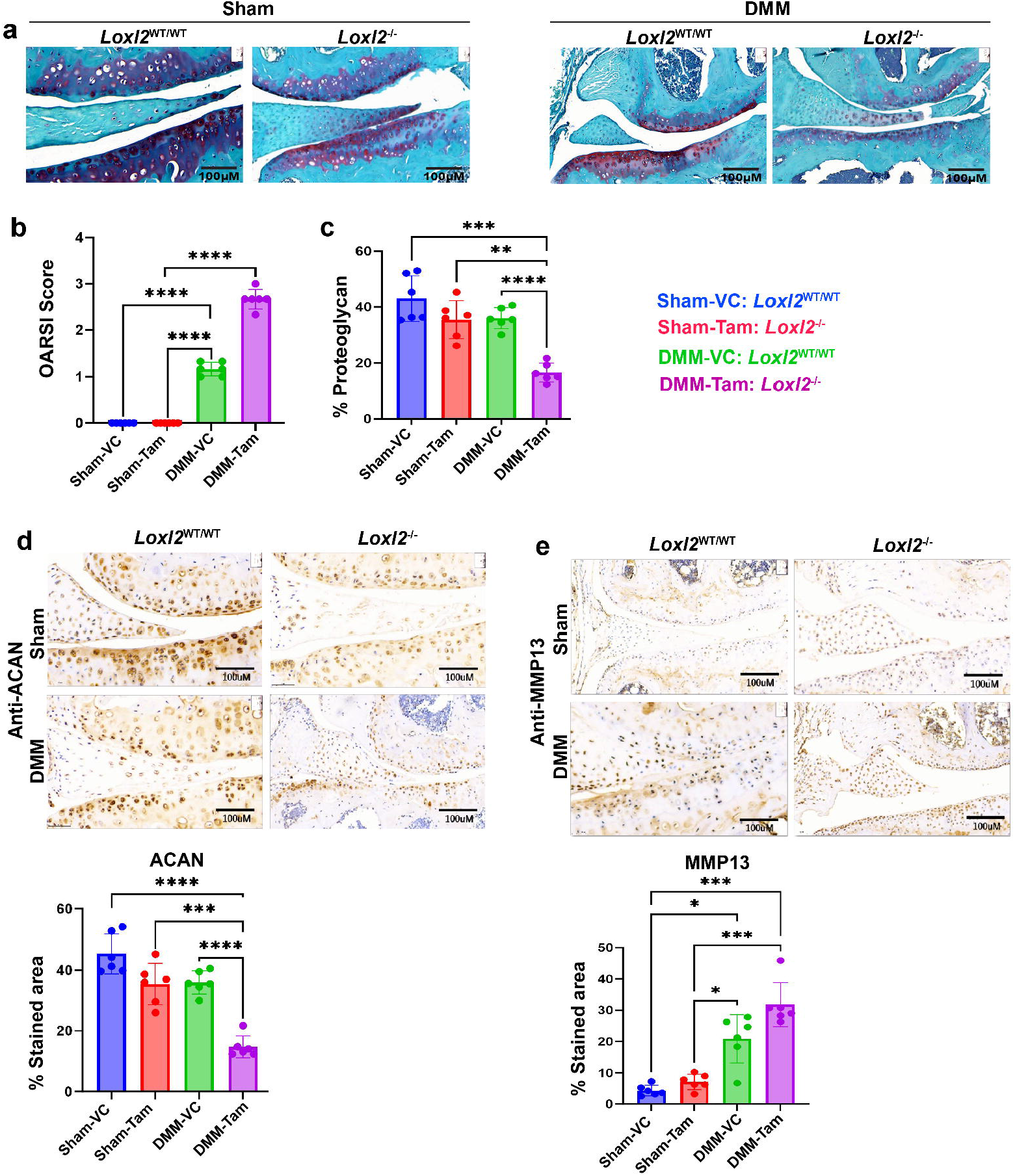
*Loxl2* deleted DMM surgery mice possess OA-pathology. **a)** Safranin-O staining of *Loxl2* wildtype and *Loxl2* deleted mice knee joint after DMM and sham surgery. **b)** OARSI score showing OA pathological phenotype with DMM surgery and *Loxl2* deletion enhancing the OA severity as represented by the elevated OARSI score. c) Bar plots showing reduced proteoglycans (GAG) deposition in DMM *Loxl2* deleted mice knee joint. d) Reduced aggrecan levels in *Loxl2* deleted sham and DMM surgery models. e) Elevated levels of MMP13 in wildtype and *Loxl2* deleted DMM surgery models. *p<0.05, **p<0.01, ***p<0.001, ****p<0.0001. P values were calculated using Brown-Forsythe and Welch’s Analysis of Variance (ANOVA) test.

### Transcriptomics analysis confirms the increased knee OA severity with *Loxl2* deletion

*Loxl2* deletion decreased in the mRNA levels of *Loxl2* and *Acan*, whereas increased *Mmp13* (Fig. 3a). To evaluate degenerative and inflammatory changes in the cartilage, RNA-seq was performed to determine the status of catabolic processes, macrophage levels, pro-inflammatory signaling, and pain perception after *Loxl2* deletion in PTOA mice. Eight mouse samples were subjected to RNA-Seq (n=4/condition). Differential gene expression (DGE) analysis detected approximately 4046 dysregulated genes (adjusted p-value <0.05) with *Loxl2* deletion (Fig. 3b; Table S1). Interestingly, key OA markers, such as Mmp13, Postn, and Col1a1, were highly upregulated (Fig. 3c), suggesting that *Loxl2* deletion in DMM mice favors OA-like molecular changes. GSEA was performed to identify affected gene sets and pathways. As expected, in *Loxl2* deleted PTOA mice, collagen and ECM degradation pathways were highly enriched, with increased gene sets related to collagen catabolic processes (Fig. 3d). Immune activation and inflammation-related pathways were also significantly enriched, including IL-1β and IL6 production, mast cell and macrophage activation, cytokine production, and the sensory perception of pain (Fig. 3d). Furthermore, we observed a significant increase in the expression of the OA-associated genes. Dysregulated genes in *Loxl2* deleted PTOA mice were significantly correlated with 134 previously reported human knee OA-associated genes from clinical samples (Fig. 3e, Table S2) ^17,18^. Overall, *Loxl2* deleted gene signatures suggest the activation of OA-specific gene sets and pathways and a positive correlation with human knee OA physiology.

**Fig. 3:**
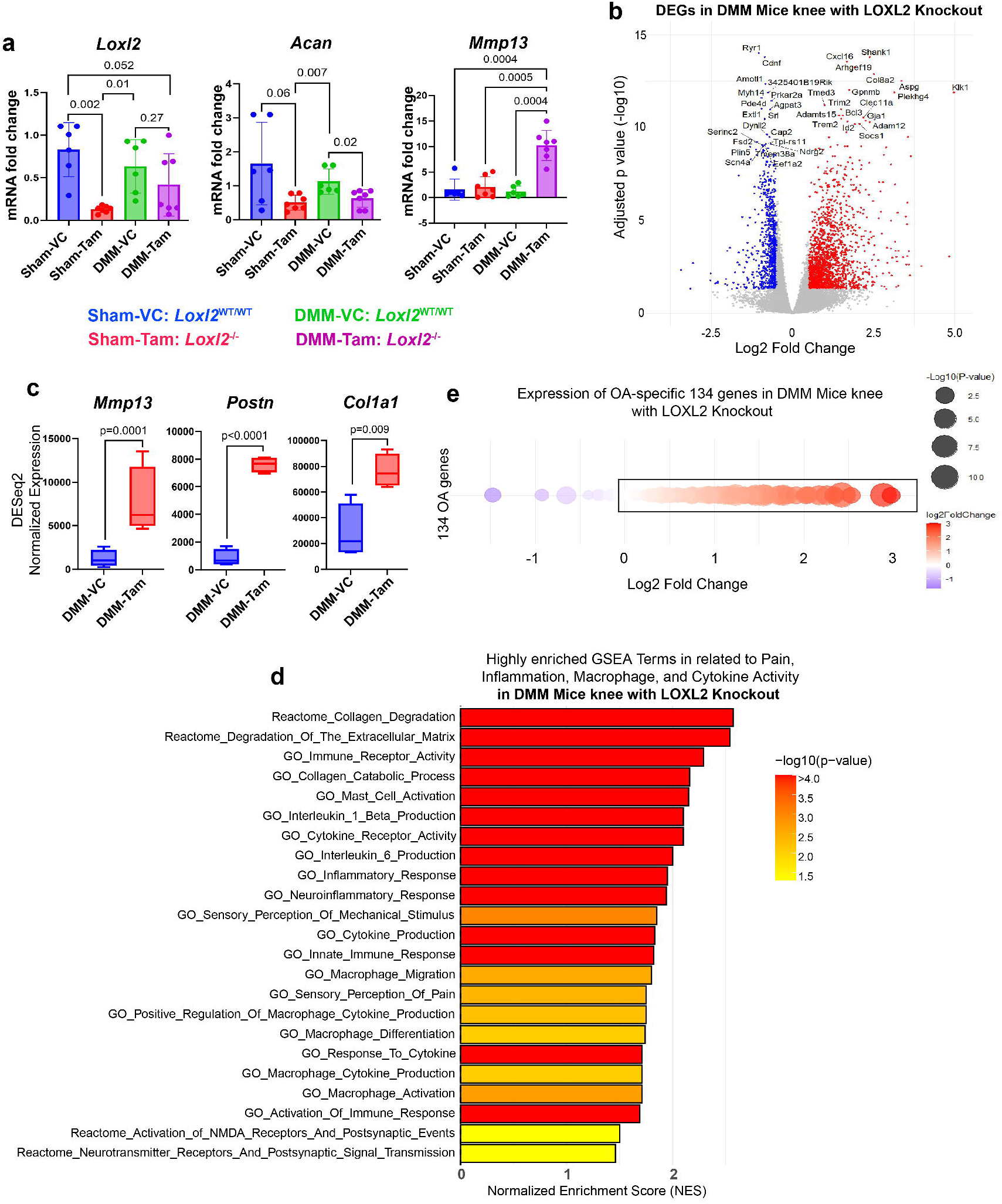
Transcriptomic analysis confirms the increased knee OA severity with *Loxl2* deletion in DMM induced knee-OA. **a)** RT-qPCR analysis showing reduced mRNA expression of *Loxl2 and Acan,* in *Loxl2* deleted DMM and sham surgery models, whereas elevated mRNA expression of *Mmp13* in *Loxl2* deleted DMM surgery mice; P values were calculated using Brown-Forsythe and Welch’s ANOVA test. b) Volcano plot of total differentially expressed genes (DEGs) in *Loxl2* deleted DMM mice compared to *Loxl2* intact DMM mice using DESeq2. Top up- and downregulated genes are labeled. Here, upregulated genes are highlighted in red and downregulated genes are highlighted in blue with significant adjusted p-value cutoff |<0.05| (Benjamini-Hochberg method). c) The expression of key OA marker genes *Mmp13, Postn*, and *Col1a1* in *Loxl2* deleted DMM mice; p values were calculated with Wald test in DESeq2. d) GSEA analysis of top DEGs from *Loxl2* deleted DMM mice has higher enrichment of pain, inflammation, macrophage activation and cytokine activity gene sets. Here, we performed gene ontology biological processes (GOBP) and Reactome pathway analysis. e) Bubble plot showing increased expression of 134 OA specific pathogenic transcription factors in *Loxl2* knockout DMM mice, which were identified previously from bulk RNA-seq of human knee-OA patients.

### Human OA promotes collagen degradation, inflammation, and pain-related gene network similar to DMM LOXL2 knockout mice

To evaluate the specific pathways and networks affected in OA, human knee OA bulk RNA-seq data were reanalyzed using publicly available profiles of 18 healthy and 20 knee OA samples^17^. Differential expression analysis was performed to screen for the dysregulated genes associated with OA (Fig. 4a, Table S3). Overexpression of the OA-specific genes MMP13, COL1A1, and POSTN, and pain marker genes AIF1, TACR1, and CSF1R (Fig. 4b). OA gene networks were highly enriched in the GSEA analysis (Fig. 4c). Specifically, gene ontology and reactome pathway analyses identified collagen and ECM degradation, inflammation, IL1 and IL6 production, and pain-related gene networks (Fig. 4c). Interestingly, these gene networks were upregulated in *Loxl2* deleted DMM mice, suggesting a molecular resemblance between the human knee OA and PTOA mouse models. Therefore, our *Loxl2* deleted PTOA mouse model could serve as an OA-specific animal model to develop effective therapeutic strategies.

**Fig. 4:**
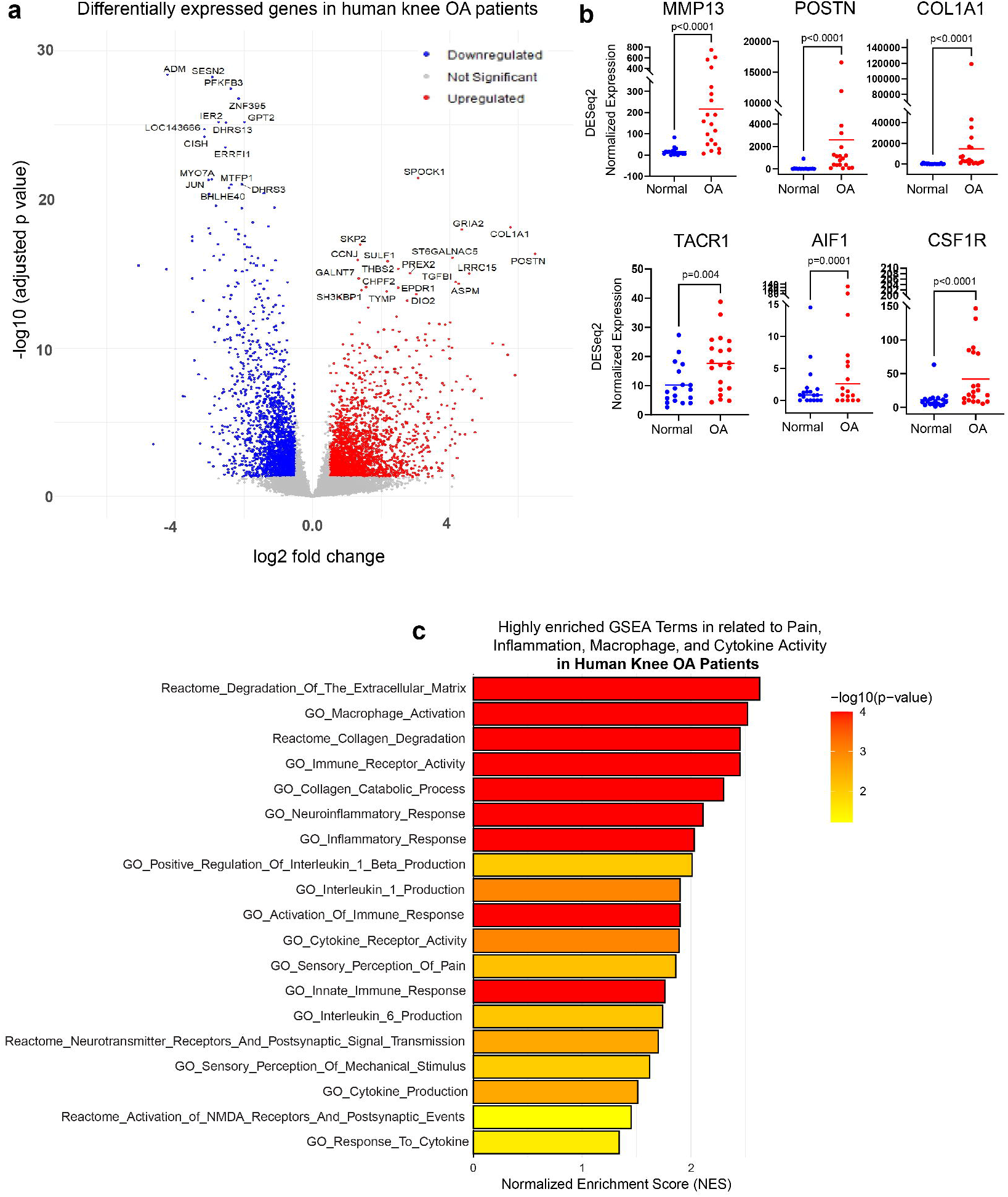
Human OA promotes collagen degradation, inflammation and pain-related gene network, similar to Loxl2 DMM knee-OA. **a)** Volcano plot shows the dysregulated genes in human knee OA patient samples. b) Expression of OA and pain-specific genes in human knee OA patients; p values were generated using the default Wald test in DESeq2. c) GSEA plot shows elevated gene networks related to cartilage/ECM degradation, inflammation, and pain.

### Loss of LOXL2 promotes synovitis and inflammatory immune cell accumulation

As indicated by the GSEA analysis, processes such as macrophage activation, differentiation, migration, and cytokine production were highly enriched when *Loxl2* was deleted. We performed CIBERSORT immune deconvolution of DEGs from Loxl2 deleted PTOA mice to further study macrophage-associated immune activity using TIMER2. We observed an increase in the proportion of macrophages in *Loxl2* deleted samples (Fig. 5a, b). We also analyzed the status of inflammatory macrophages (M1 macrophages) and found that the levels of *Cd86, Tnf, Tlr2,* and *Tlr4* were significantly increased, suggesting elevated inflammatory immune activation in the absence of LOXL2 (Fig. 5c).

**Fig. 5:**
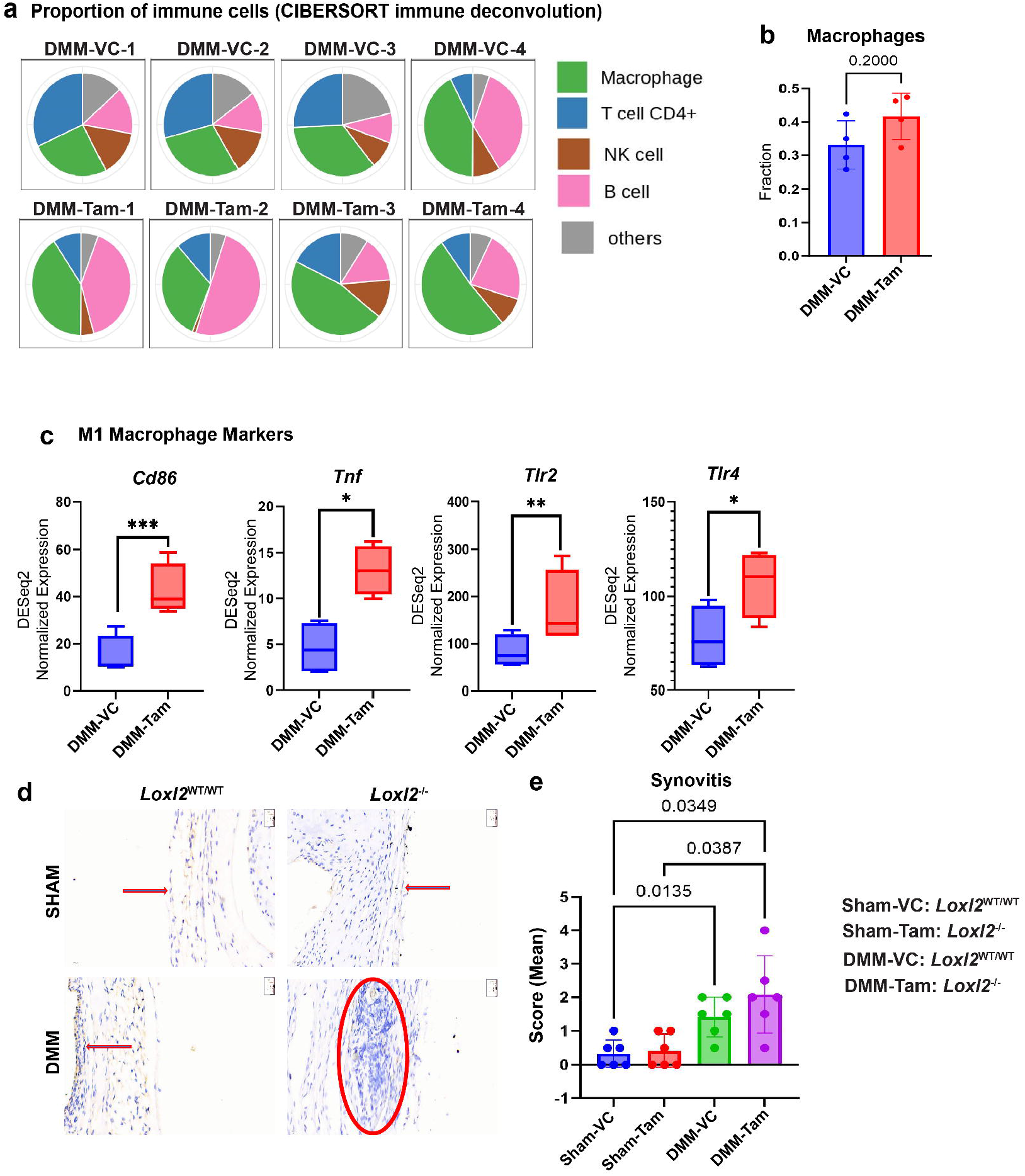
*Loxl2* deletion elevates inflammatory macrophages, induced cytokine activity, and develops synovitis. **a-b)** Immune Deconvolution with CIBERSORT algorithm in TIMER2 webtool showing enhancement of macrophage populations in *Loxl2* deleted DMM mice in all four replicates. c) Boxplots for DESeq2 normalized elevated expression of inflammatory M1 macrophage marker genes in *Loxl2* deleted DMM mice; P values were calculated by the default Wald test in DESeq2. d) H&E staining showing increased infiltration of immune cell population (blue) in synovial tissue of *Loxl2* deleted DMM surgery mice. e) Quantification of synovitis score (mean) highlights enhanced immune activity and synovitis in the *Loxl2* deleted DMM surgery mice; P values were calculated using Brown-Forsythe and Welch’s ANOVA test. *p<0.05, **p<0.01, ***p<0.001, ****p<0.0001.

The synovium is a type of connective tissue that lines the joints, tendons, bursae, and fat pads, closes the synovial space, and regulates the fluid composition. It produces lubricin and hyaluronic acid and helps in chondrocyte nourishment via synovial fluid because cartilage lacks a vascular supply. Macrophage infiltration into the synovium is prevalent in OA patients. Synovial macrophages are responsible for a large proportion of the innate immune activation and cytokine production in OA joints. The infiltration of leukocytes, primarily macrophages, is stimulated by cytokines and adhesion molecules in the synovium of OA patients ^19^. Here, *Loxl2* deleted mice subjected to DMM surgery showed synovial membrane thickening and synovitis-like changes. Immune cell infiltration was significantly higher in *Loxl2* knockout DMM mice (Fig. 5d). Synovitis was more severe in the knee joints of *Loxl2* deleted DMM mice (Fig. 5e). Synovial inflammation enhances the reactivity of peripheral nociceptive neurons, resulting in improved pain sensitivity ^20^. This suggests that LOXL2 prevents synovial inflammation and pain, and potentially prevents immune cell influx.

### LOXL2 IA injection reverses OA-related pain in DMM mice

GSEA (Reactome and GOBP) of *Loxl2* deleted PTOA mice showed that terms such as pain and neuroinflammation, sensory perception of pain, neuroinflammatory response, and activation of N-methyl-D-aspartate (NMDA) and neurotransmitter receptors were significantly enriched (Fig. 4c). Classical M1 macrophage markers, such as *Cd86, Tnf, Tlr2,* and *Tlr4* were highly upregulated (Fig. 5c).

Hence, we analyzed the expression of inflammatory pain inducer genes such as Piezo2, Aif1, Tacr1, Ptgs2, P2rx7, and Csf1r, and found that they were significantly upregulated in *Loxl2-deleted* PTOA mice (Fig. 6a). These data indicated that *Loxl2* deletion promotes pain perception in mouse knee OA.

**Fig. 6:**
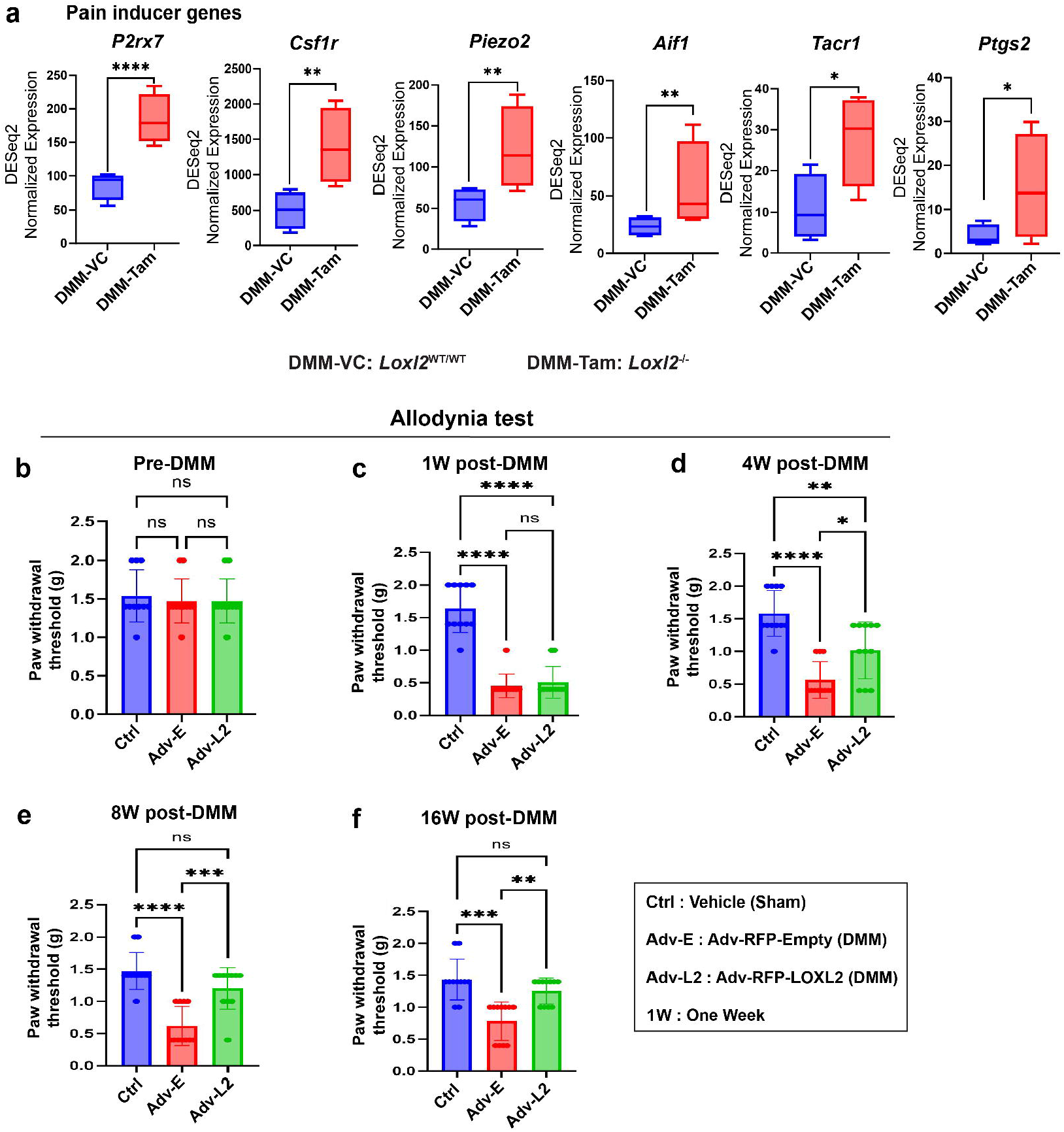
LOXL2 treatment reduces inflammation and pain in PTOA mice. **a)** Boxplots for DESeq2 normalized enhanced expression of classical pain inducer genes of knee OA in *Loxl2* deleted DMM mice models; P values were calculated by the default Wald test in DESeq2. b) Bar plot representation of unchanged mechanical allodynia response (using von Frey filaments) in untreated control, adenoviral empty vector control, and adenoviral LOXL2 overexpression mice. c) Allodynia response 1-week post-DMM; d) 4-weeks post-DMM; e) 8-weeks post-DMM; f) 16-weeks post-DMM; showing gradual recovery from allodynia pain in adenoviral LOXL2 treated DMM surgery mice; P values were calculated using Brown-Forsythe and Welch’s ANOVA test. *p<0.05, **p<0.01, ***p<0.001, ****p<0.0001, ns: p>0.05.

To test whether LOXL2 induces an anabolic response and restores knee joint function, we compared DMM mice injected with Adv-RFP-LOXL2 (LOXL2 treated) and Adv-RFP-Empty (empty vector). Both male and female C57BL/6J mice (n=10-11/condition) were injected 7 days post-DMM and subjected to a mechanical allodynia test using von Frey filaments at 1, 4, 8, and 16 weeks. The pre-DMM surgery group showed an unaltered paw withdrawal response (Fig. 6b); however, the post-DMM surgery group responded differently depending on the treatment and duration post-IA injection. One week post-DMM, there was an adverse paw withdrawal response in the empty vector and LOXL2-treated group, suggesting an elevated pain sensation sustained for a week (Fig. 6c). Interestingly, four weeks post-IA injection, recovery was observed in the LOXL2 treated group compared to that in the empty vector group (Fig. 6d). This LOXL2 mediated recovery persisted for 8 and 16 weeks, respectively (Fig. 6e, f). Overall, LOXL2 therapy in PTOA mice reduced pain and discomfort in the OA knee by promoting the restoration of normal cellular and mechanical homeostasis.

### LOXL2 IA injection protects against DMM-induced progressive changes

Next, we performed a mouse treadmill exhaustion test to assess the effects of exercise intensity on knee joints, leading to exhaustion. The pre-DMM surgery models treated with empty vectors or LOXL2, along with the untreated control, did not show any variation in the running distance (Fig. 7a) or time (Fig. 7c). In the post-DMM surgery group, empty vector-treated mice covered less distance than untreated controls one month after IA injection (Fig. 7b). These mice ran faster than untreated control mice (Fig. 7d). As expected, running distance and time were restored to normal one month after LOXL2 IA injections (Fig. 7b, d).

**Fig. 7:**
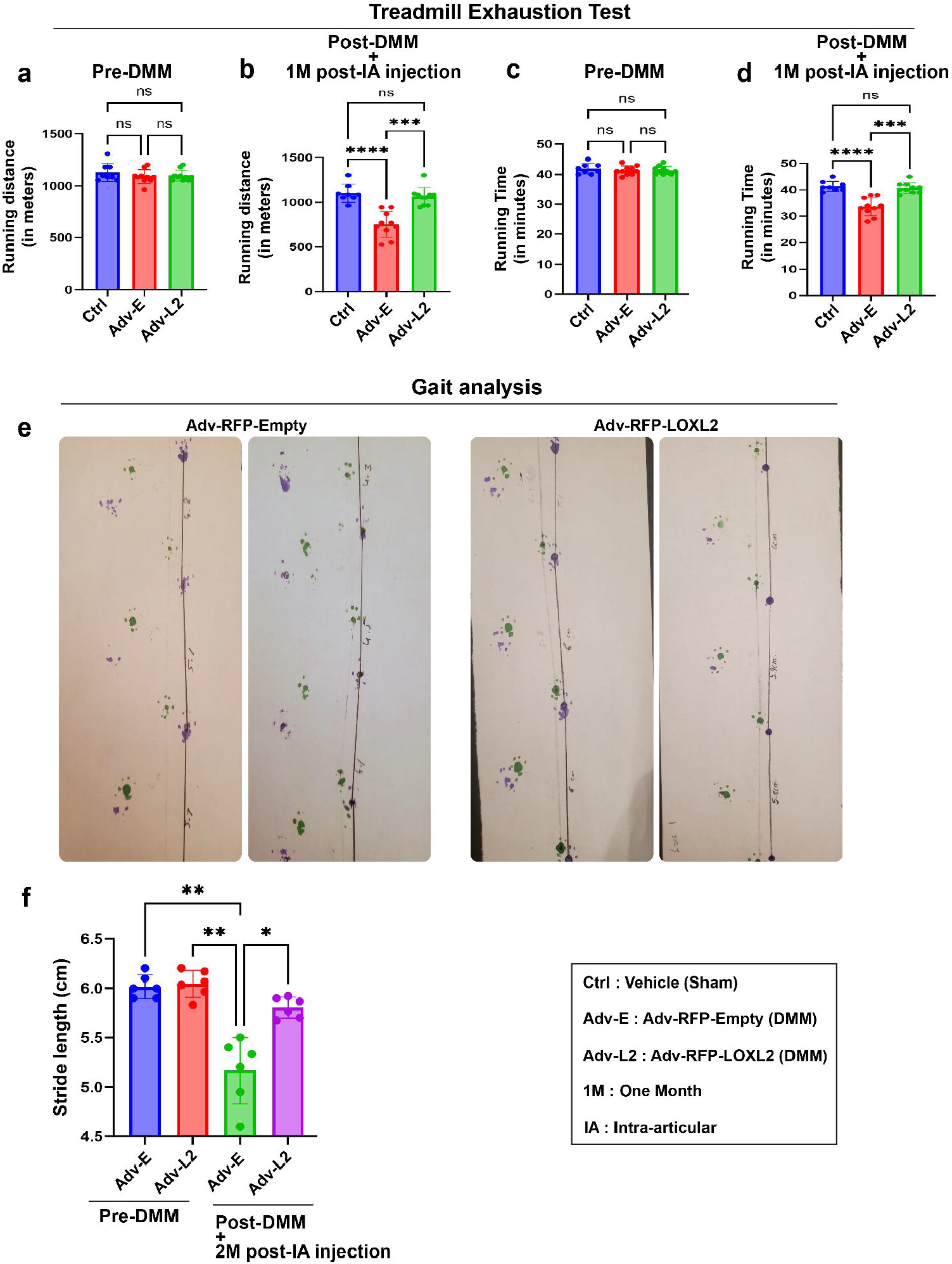
LOXL2 IA injection protects against DMM-induced progressive changes. **a)** The treadmill exhaustion test found an unchanged running distance in all three mice groups before DMM surgery. b) Recovery in running distance 1-month after Adenoviral LOXL2 treatment in DMM surgery mice. **c)** The treadmill exhaustion test showed an unchanged running time in all three mice groups before DMM surgery. **d)** Recovery in running time 1-month after adenoviral LOXL2 treatment in DMM surgery mice. **e)** Stride length measurement of DMM hind limb. **f)** Bar plot representation of Gait analysis of pre- and post-DMM surgery mice groups after adenoviral LOXL2 treatment. *p<0.05, **p<0.01, ***p<0.001, ****p<0.0001, ns: p>0.05. All P values were calculated using Brown-Forsythe and Welch’s ANOVA test.

To minimize pain and load on the damaged knee joint, mice with knee OA often walk with reduced weight-bearing on the affected leg, which may appear as a subtle limp, and exhibit altered stride length measured using gait analysis techniques to gauge the severity of OA and efficacy of treatments ^21,22^. We performed gait analysis using six mice per condition and measured stride length in the pre-DMM and post-DMM surgery groups after two months of empty vector and LOXL2 treatment (Fig. 7e). The pre-DMM group did not show any measurable variation in stride length; however, the post-DMM empty vector group had a significantly reduced stride length, which was successfully recovered in the LOXL2-treated group, suggesting reduced pain and near-normal recovery in the LOXL2-treated PTOA mice (Fig. 7f). Overall, LOXL2 treatment increased endurance and reduced pain, leading to successful recovery from structural and functional damage in the PTOA mice. This LOXL2 gene therapy can be further explored as a potential candidate for OA treatment.

## Discussion

The overarching goal of this study was to determine whether LOXL2 loss promotes OA and pain, whereas LOXL2 gain reverses OA-related pain and joint function. This potential is also supported by LOXL2’s combined chondroprotective and anabolic effects in OA ^5^. Our study demonstrated that LOXL2 depletion significantly enhanced OA severity by promoting cartilage degradation, inflammation, and pain in the knee joints of mice and its overexpression alleviated these effects.

Mechanistically, IHC analysis of LOXL2 knockout mouse knee cartilage showed a drastic reduction in the levels of aggrecan and a significant increase in the levels of MMP13. This was further supported by our DMM surgery-induced *Loxl2* knockout models, where RNA sequencing followed by immune cell deconvolution and gene set enrichment analysis showed elevated levels of inflammatory macrophages, hyperactivation of cytokine signaling, and induction of hyperpain sensitivity pathways. LOXL2 depletion also favors chondrocyte dedifferentiation and promotes synovitis in the knee joint. Furthermore, overexpression of LOXL2 using an adenovirus delivery system in the DMM-induced mouse knee alleviated mechanical allodynia outcomes during the course of four–16 weeks post-IA injections. This overexpression also increased endurance and resulted in better limping outcomes as observed in the treadmill test and gait analysis.

LOXL2 has both intracellular and extracellular functions, as it localizes to the nucleus, perinuclear region, cytoplasm, and extracellular matrix (ECM) ^7,23^. The conventional extracellular role of LOXL2 is to promote collagen cross-linking and maturation. LOXL2-induced collagen crosslinking enhances the tensile strength of articular cartilage and resistance to collagen proteolysis ^7^. ADAMTS4/5 and MMP13 are major proteinases that degrade proteoglycans and collagens, respectively, leading to pathological changes in the OA cartilage. RNA-seq followed by DGE analysis of *Loxl2* deleted DMM mice revealed hyperexpression of MMP13 and ADAMTS4/5 and higher enrichment of collagen and ECM degradation gene sets, suggesting an unconventional role of LOXL2 in protecting the knee joint from degenerative OA and maintaining healthy chondrocyte homeostasis.

Macrophage infiltration, followed by the release of pro-inflammatory cytokines due to cartilage damage in the knee joint, is a major event that drives inflammation and joint pain in patients with OA, aggravating disease severity. Activated macrophages produce inflammatory mediators such as interleukin-1 beta (IL-1β) and tumor necrosis factor-alpha (TNF-α), which further damage cartilage tissue. M1 polarized proinflammatory macrophages contribute to cartilage destruction and inflammation ^24–26^. Our DGE analysis, followed by CIBERSORT immune deconvolution, detected an increased number of macrophages in DMM-induced *Loxl2* knockout mice. Overexpression of key M1 macrophage marker genes, such as *Cd86, Tnf,* and *Tlr2/4* was also detected, suggesting their increased levels in the absence of LOXL2. Gene set enrichment analysis revealed the upregulation of gene sets related to macrophage activation, differentiation, and migration. Furthermore, the mast cell activation gene set was highly enriched, which could lead to the release of histamines and cytokines, eventually worsening the disease severity due to hyperinflammatory activity. They also release enzymes, such as tryptase, which can directly break down cartilage and extracellular matrix components ^27^. As M1 macrophages and mast cell activation are the major outcomes of *Loxl2* deletion in the cartilage, cytokine activity (production and activation) is expected to increase and contribute to inflammation. Interestingly, our study detected higher enrichment of cytokine production and cytokine receptor activity gene sets, contributing to the innate immune response.

The increased expression of pain-related genes, such as *Piezo2, Aif1, Tacr1, Ptgs2, P2rx7,* and *Csf1r,* in our mouse knee OA DMM model after *Loxl2* deletion resulted in considerable alterations in pain-signaling pathways. These findings shed light on the mechanisms driving pain sensitization in OA as well as the specific roles that LOXL2 plays in regulating these processes. PIEZO2 encodes an important mechanosensitive ion channel involved in nociception and mechanotransduction. The greater pain observed in OA models is consistent with its overexpression, which suggests higher sensitivity to mechanical stress inside the joint ^28,29^. Another factor contributing to inflammation-associated pain is AIF1 (Allograft Inflammatory Factor 1), a marker of activated macrophages and microglia, which suggests an increased neuroimmune inflammatory response ^30^. TACR1 encodes the neurokinin 1 receptor and transmits pain via substance P, a neuropeptide associated with chronic pain ^31^. Elevated TACR1 expression highlights its potential role in neurogenic inflammation in the absence of LOXL2. Furthermore, overexpression of PTGS2 (prostaglandin-endoperoxide synthase 2, or COX-2) indicates an elevated prostaglandin pathway, which is a well-known contributor to inflammatory pain ^32^. The increased expression of P2RX7, a purinergic receptor involved in ATP-mediated pain signaling and inflammasome activation, suggests a more significant purinergic contribution to nociceptive pathways and neuropathic pain ^33,34^. Finally, elevation of CSF1R (a colony-stimulating factor 1 receptor) indicates enhanced macrophage proliferation and activation, which is likely to exacerbate the inflammatory environment and neuropathic pain within the joint ^35^. These data indicated that LOXL2 may play a protective role in modifying pain pathways in knee OA by reducing pro-nociceptive and inflammatory pain mediators.

Intra-articular gene transfer by adeno-associated viruses (AAVs) is advantageous in clinical trials ^36^ and could be a delivery strategy for LOXL2 therapies. In our study, intra-articular injections of AAV-LOXL2 into the knees of pre- and post-DMM mice resulted in variable mechanical allodynia. Four to 16 weeks after LOXL2 injection, the allodynia response tended to recover in DMM mice compared to that in the empty vector-injected controls. This finding aligns with our LOXL2 knockout DMM model in which LOXL2 depletion seemed to aggravate inflammatory pain. However, overexpression assists in pain relief, resulting in improved allodynia outcomes. Overexpression of LOXL2 also enhanced the endurance of DMM mice one month after IA injection, as determined by the treadmill test. Additionally, LOXL2 helped recover from the shortened stride length observed in the post-DMM empty vector controls, indicating less pain and nearly normal recovery in knee OA mouse models.

Our *Loxl2* knockout mouse models, which are representative of the knee OA model for preclinical small-animal *in vivo* studies, showed a strong positive correlation with 134 OA transcription factors identified from bulk RNA sequencing analysis of cartilage from patients with knee OA ^17,18^.

In conclusion, LOXL2 is critical for maintaining articular cartilage, ECM, ACAN, and knee joint functions. Co-relative analysis of the human OA-related pathogenic signature with structural and functional studies performed in *Loxl2* knockout mice showed that LOXL2 may function as a fundamental regulator of joint tissue homeostasis, influencing both baseline and stress-induced transcriptional networks. The loss of LOXL2 is a “predisposing factor” that initiates a vicious cycle of progressive degenerative changes and pain. LOXL2 exhibits chondroprotective and chondroregenerative properties by restoring chondrocyte signaling, ECM, collagen crosslinking, and other functional properties, thereby alleviating pain. Furthermore, loss of LOXL2 in mice indicates the natural prognosis of OA-like changes and could be used as a model to study OA.

## Materials and Methods

### Ethical Compliance

Approval was obtained from the Boston University Institutional Animal Care and Use Committee (IACUC; approval number: AN-15387). Animal studies conformed to the ARRIVE guidelines.

### Cartilage-specific *Loxl2*knockout mice

*Loxl2* floxed (fl) mice were obtained from Dr. Cano’s laboratory in Madrid, Spain ^13^. We used the *Agc1^tm(IRES-CreERT2)^* murine line (Jackson Laboratories, #019148) generated by Henry et al. ^14^, which has a similar *Acan* promoter/enhancer expression pattern as another murine line ^37^. To avoid any impact on cartilage development, we used Tm-inducible ER^T2^ to knock out *Loxl2* in cartilage by crossing the Acan-CreERT2 strain ^15,16^ with *Loxl2*^fl/fl^ mice, in which exon 2 was flanked by loxP sites [13], to generate Tm-inducible Acan-Cre^ERT2^;*Loxl2*^fl/fl^ mice. We optimized the protocol for mice, which includes one 100-μL intraperitoneal injection/day (75 mg/kg in corn oil) for five consecutive days ^38,39^, followed by a single maintenance injection every month.

### Establishment of DMM

The DMM protocol involves cutting the ligament to the lateral side (instead of the midpoint, as in earlier protocols) using a dissection microscope. The knee joint was rotated medially and bent at 45°, followed by an incision of the knee joint. The medial meniscus was cut laterally by using a blade 11 curved/straight blade. Successful DMM surgery revealed visible femur and tibia at the knee junction, and the medial meniscus was transected and popped out under higher magnification. Finally, the joint capsule was sutured using Vicryl suture 4.0, followed by skin suturing. Thus, we established a modified DMM model with the advantages of a minimal incision size, reproducibility, and ease of validation.

### Histology and immunostaining

Knee joints from mice were paraffin-embedded, decalcified, and subjected to histological analysis and immunostaining. Safranin-O/Fast Green (American Mastertek Inc.) staining was performed as previously described ^5^. OARSI scoring was performed according to recommendations. Three sections from four mice per group were deparaffinized, immunostained with specific antibodies to detect LOXL2, ACAN, and MMP13 (Abcam), and visualized with HRP-linked anti-rabbit antibodies. Stained tissues were scanned using a digital slide scanner (Panoramic MIDI, 3D Histech).

### Treadmill exhaustion test

Treadmill behavior is helpful in preclinical behavioral assessments of chronic pain and inflammation ^40^. Treadmill analysis was performed using a standard protocol ^41^ before and after the DMM surgery. Mice were acclimatized to treadmill running (TSE Systems) for three consecutive days, followed by resting for 1 d before performance evaluation ^41^. Acclimatization consisted of a 5-minute rest on the treadmill conveyor belt, followed by 5 min of running at 7.2 m/s and 5 min at 9.6 m/s. On day 0, the mice were subjected to a graded maximal running test consisting of an initial 5-min rest, after which the running protocol commenced at 4.8 m/min, gradually increasing by 2.4 m/min every 2 min. At all times, the belt was kept at a 5-degree incline. Maximal running speed was defined as the fastest speed at which mice could run for five consecutive seconds without touching the electric shock grid at the back of the treadmill. Here, exhaustion was defined as the time when the mice were unable to avoid an electrical shock grid at the back of the treadmill for 3 consecutive seconds or 45 min, whichever came first.

### Allodynia test (von Frey Test for Nociception Assessment)

Allodynia was evaluated using the von Frey–Hairs test by pricking the hind paw to 3-5 times with filaments of different sizes. When measuring the withdrawal reflex in response to filament touch, mice with pain were more sensitive to smaller filament sizes. The researcher conducting the test was blinded to the experimental groups.

### Gait analysis

All mice were acclimatized to walking through a tunnel measuring 2.5, 3, and 13 cm in width, height, and length, respectively. We fabricated an end chamber measuring 4 inches in length and width and 3 inches in height into which the tunnel led. The tunnel was placed on white paper, onto which the mouse limb prints were recorded. It was 10 cm wide and 60 cm long. The hind paws of the mice were painted with inks of different colors to differentiate them from the operated and unoperated knees. Mice were allowed to walk to a tunnel with their paws imprinted on a white piece of paper. A complete walking cycle (right-to-right-distance), which is referred to as “relative step length” of 4. Additionally, footprint analysis was used to study paw posture and parameters such as toe spread (TS), intermediate toe spread (ITS), and print length (PL) ^42^.

### RNA-sequencing and bioinformatics analyses

Total TRIzol reagent was extracted from all samples (n=4/condition), which were then sent for RNA sequencing at Novogene in Sacramento, California. Raw reads were quality-controlled before being clipped or pre-processed using an in-house Novogene Perl script. was mapped to the Mus musculus reference genome (mm10). Feature Counts (v1.5.0-p3) were used to quantify data. Differential Gene Expression (DGE) analysis was performed using DESeq2 R/Bioconductor. Biological processes and reactome pathways were identified using gene set enrichment analysis (GSEA) (v4.3.2). Volcano and GSEA horizontal bar plots were created using ggplot2 R package. Box/bar plots were generated using GraphPad PRISM (v10.1.0).

### Reanalysis of Human Knee OA patient samples

Raw RNA-seq count data for 38 healthy human (n=18) and knee-OA (n=20) samples were acquired from the NCBI Gene Expression Omnibus (GEO) database with accession ID GSE114007. DGE analysis was performed using the DESeq2. R/Bioconductor. Downstream analysis and plotting were performed as described above.

## Acknowledgment

This study was supported by an NIH grant R01 DE031413 (MB).

## Ethical Compliance

All procedures performed in this study involving C16BL/6J mice were in accordance with the ethical standards of the institutional and/or national research committees.

## Author Contributions

Faiza Ali, Rajnikant Dilip Raut, Chumki Choudhury, Amit Kumar Chakraborty, Cheyleann Del Valle-Ponce De Leon, and Manish V. Bais performed the experiments and analyzed the data. Pushkar Mehra and Manish V. Bais helped with conception, interpretation, and manuscript editing. Manish V. Bais conceived and designed the study. Rajnikant Dilip Raut, Faiza Ali and Manish V. Bais interpreted the data and wrote the manuscript.

## Declaration of generative AI in scientific writing

The authors declare that they have not used AI to write or edit this manuscript.

## Data Availability

Raw RNA sequencing data for *Loxl2* deleted mice knee DMM were submitted to NCBI GEO under accession ID GSE285114 (reviewer tokens are available upon request).

---

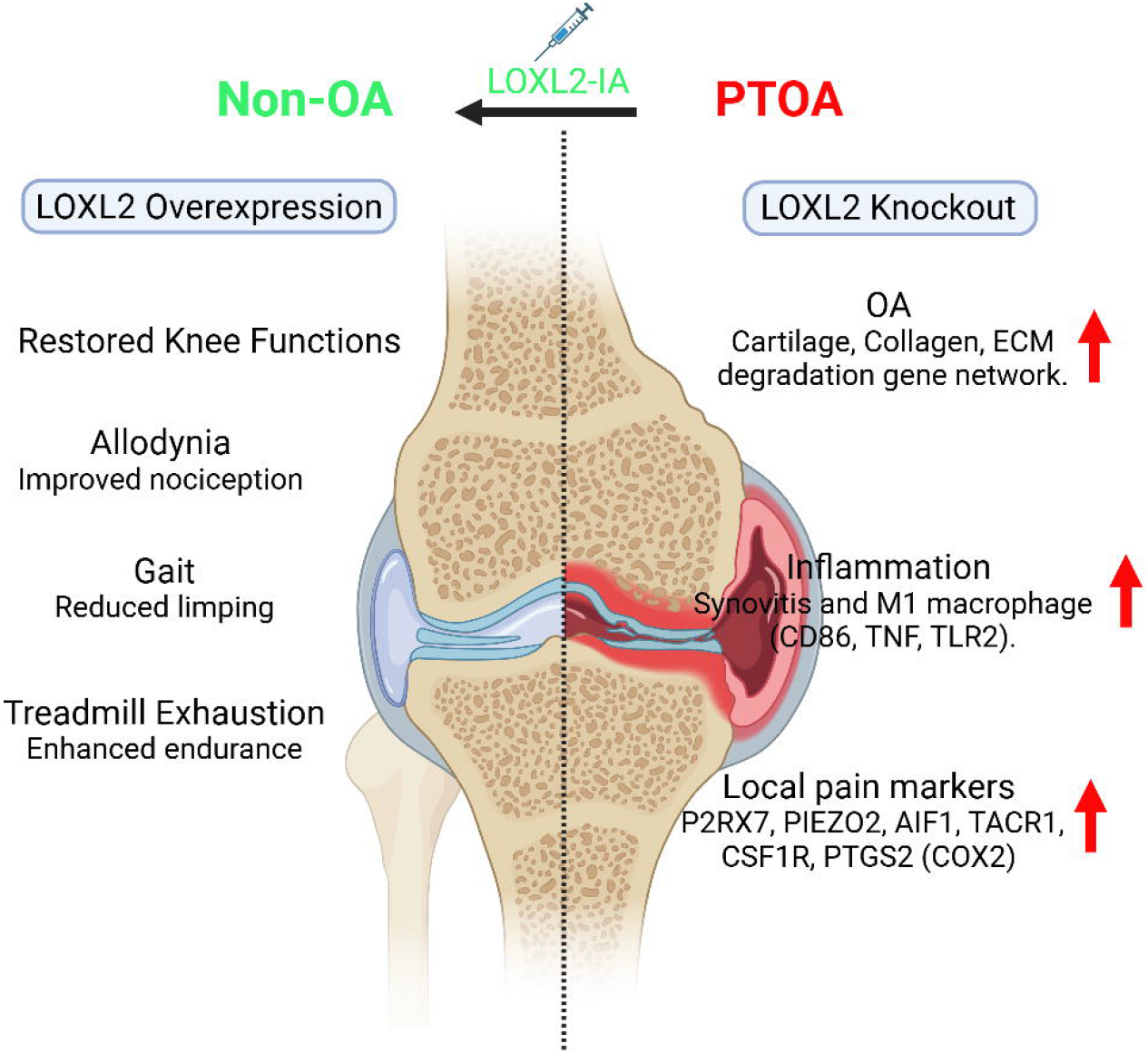

